# Dynamic Electrical Stimulation of Sites in Visual Cortex Produces Form Vision in Sighted and Blind Humans

**DOI:** 10.1101/462697

**Authors:** Michael S. Beauchamp, William Bosking, Ping Sun, Brett Foster, Soroush Niketeghad, Nader Pouratian, Daniel Yoshor

## Abstract

Visual cortical prosthetics (VCPs) offer the promise of restoring sight to blind patients. Electrical stimulation of a single site in visual cortex can reliably produce a percept of a spot of light in a fixed visual field location, known as a phosphene. Researchers developing VCPs have assumed that multiple phosphenes produced by concurrent stimulation of multiple sites in visual cortex can combine to form a coherent form, like pixels in a visual display. However, existing data do not support this assumption. Therefore, we developed a novel stimulation paradigm for VCPs termed *dynamic current steering* in which the visual form to be conveyed is traced on the surface of visual cortex by electrically stimulating electrodes in a dynamic sequence. When tested in sighted and blind subjects, this method of stimulating visual cortex allowed for the immediate recognition of a variety of letter shapes without training and with high accuracy.

**One Sentence Summary:** Stimulating human visual cortex using dynamic patterns of activity allows both blind and sighted patients to perceive visual percepts of useful forms.

## Introduction

In most patients with acquired blindness, only the eyes or optic nerves are damaged. This has inspired hope for a visual cortex prosthetic (VCP) that would pass visual information from a head-mounted camera directly into the intact visual cortex (reviewed in *1*). VCPs rely on the fact that stimulating a given site in visual cortex with electrical current can produce a percept of a small flash of light, known as a phosphene, at a specific location in the visual field. Because of the retinotopic organization of visual cortex, implanting an array of multiple electrodes at different cortical locations allows for the creation of phosphenes spanning a range of the visual field, with each stimulated electrode contributing one phosphene (*2, 3*). Modern computer engineering has advanced to the point that wirelessly powered and controlled devices containing dozens of electrodes can be implanted into visual cortex, leading to worldwide efforts to develop clinically-usable VCPs (*4-6*).

However, there is a serious stumbling block. All existing and proposed VCPs use a stimulation paradigm in which multiple electrodes are concurrently stimulated, assuming that this will produce multiple phosphenes that perceptually combine into coherent forms, analogous to pixels in a computer display. However, existing data does not support this assumption. For instance, in the clinical trials of Dobelle and colleagues in the 1970s, stimulation of multiple electrodes produced only percepts of multiple isolated phosphenes which did *not* combine into forms (*7*); similar results were reported by Schmidt and colleagues twenty years later (*8*).

One explanation for the failure of the conventional electrical stimulation paradigm is the unnatural activity that it evokes in visual cortex. When viewing natural scenes, only a relatively small fraction of neurons in early visual cortex tuned to features in the scene are active. Visual experience coupled with Hebbian learning ensures that these neurons preferentially connect to each other and to neurons downstream areas, allowing the creation of neurons that represent objects. An example of this process is visual perceptual learning of letter forms. Individual letters activate specific combinations of orientation selective neurons in early visual cortex: for instance, a visually-presented “T” would activate neurons selective for horizontal and vertically oriented lines at adjacent locations within the visual field. The combination of features identified in early visual cortex then activates neurons selective for letter identity in higher areas, especially the visual word form area (*9, 10*). Even if early visual cortex is intact, damage to higher areas results in profound impairments in the ability to recognize visual motion, letters, faces and other complex visual forms (*11-13*).

In contrast to the selective activation of tuned neurons produced by real visual stimuli, electrical stimulation activates an effectively random set neurons in the immediate region of the electrode (Histed et al 2009). Electrical activation of early visual areas (V1/V2/V3) can result in a percept of a simple phosphene (*14*), but this non-selective activation of spatially contiguous neurons in may not effectively propagate to higher visual areas to produce complex percepts as normally occurs with natural vision. This would explain prior reports that multiple electrode stimulation in V1 produced only isolated phosphenes, not forms (*7, 8*). The observation that electrical stimulation of late visual areas usually produces no percept whatsoever (*14*) suggests that a complex visual percept requires the generation of a highly complex pattern of activation in later visual areas.

To overcome the failure of the conventional paradigm of concurrent multi-electrode stimulation for a VCP, we developed a novel stimulation paradigm termed *dynamic current steering.* Figure 1 illustrates this stimulation paradigm by analogy to tracing letters on the palm, the technique used to teach Helen Keller, the pioneering advocate for the blind, to communicate (*15*). To convey the letter “Z” though touch, one could press multiple probes arranged in a “Z” pattern into the palm (Fig. 1). However, this produces an incoherent percept of a touch without coherent form. Alternatively, one could press a single probe into the palm and move it in a trajectory that matches the “Z” shape (Fig. 1B), which immediately produces a letter percept.

**Figure 1.**
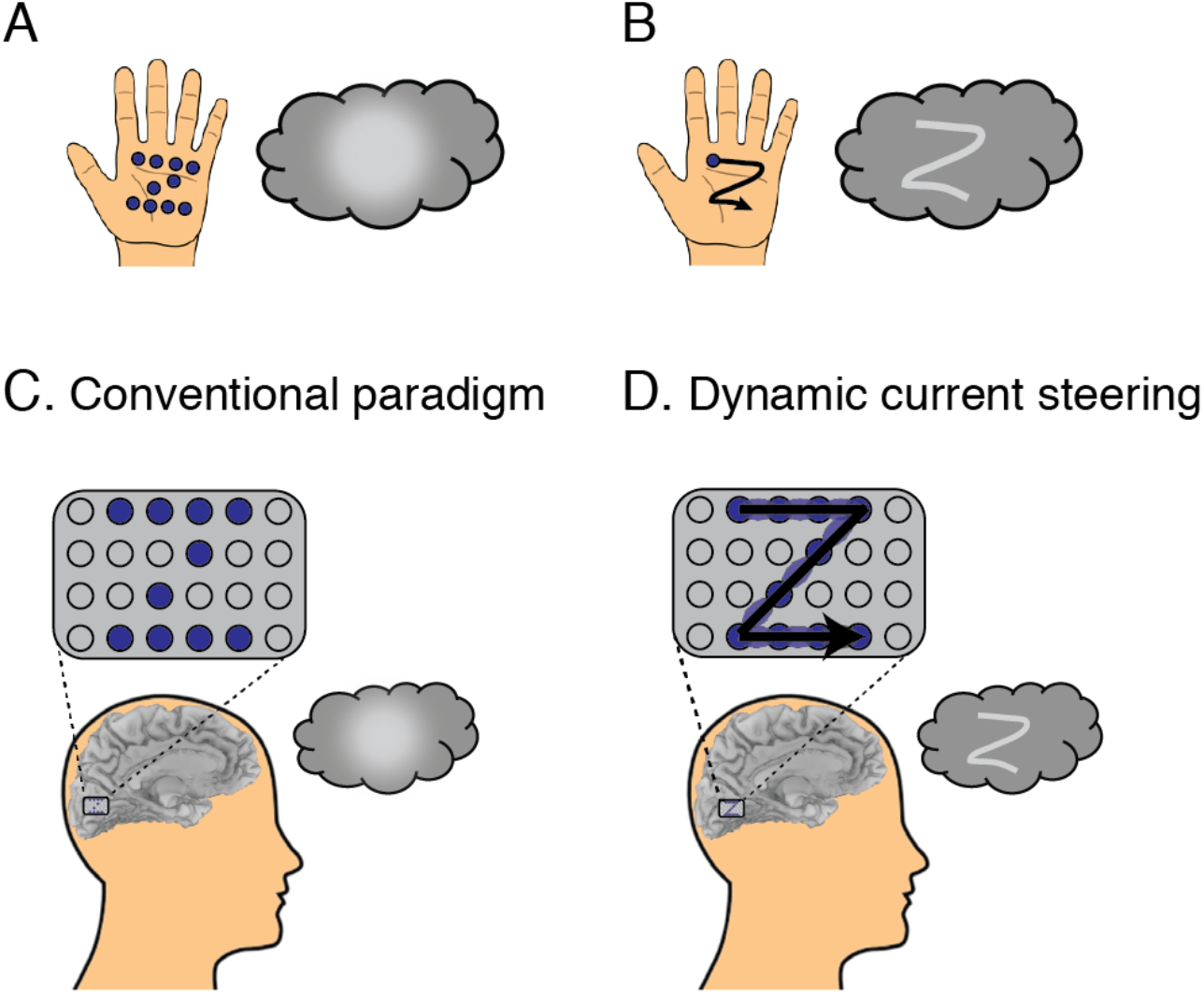
Stimulation paradigms for cortical visual prosthetics. **(A)** To convey a letter through touch, one could press multiple objects (blue dots) into the palm of the hand, forming the shape of the letter. However, this results in an amorphous percept (gray blob in thought bubble). **(B)** Alternately, one could trace the shape of the letter using a single probe (single blue dot) traced across the palm in a trajectory (black line with arrow) that matches the desired shape, producing a letter percept (“Z” shape in thought bubble). **(C)** In a cortical visual prosthetic, an electrode grid (gray rectangle) is implanted over visual cortex. With the conventional static stimulation paradigm, electrical stimulation is delivered concurrently to some electrodes (blue circles) but not others (gray circles), resulting in an amorphous percept. **(D)** In the dynamic current steering paradigm, electrical stimulation is delivered to electrodes in temporal sequence, in a trajectory that matches the desired shape. To increase the fidelity of the trajectory, current steering is used to create virtual electrodes (light blue ellipses) by stimulating adjacent electrodes simultaneously. This results in the percept of a shape matching the trajectory (“Z”).

The conventional VCP paradigm is similar to the multiple probe approach: multiple electrodes are stimulated at once, resulting in an incoherent percept without a clear form (Fig. 1C). The dynamic current steering paradigm is analogous to tracing letters on the palm, except that the trajectory of the desired letter is formed on the surface of the cortex by electrically stimulating electrodes in a dynamic sequence (Fig. 1D). The resolution of the trajectory is limited by the density of the implanted electrodes, but resolution can be enhanced with current steering: if current is passed through two adjacent electrodes, a virtual electrode is created midway between them (*16*). Combining dynamic stimulation with current steering allows a smooth trajectory of the desired form to be created, resulting in a coherent percept of the desired form.

## Results

To explore the utility of dynamic current steering in visual cortex, we studied sighted patients temporarily implanted with intracranial electrodes as part of clinical monitoring for epilepsy surgery. Novel arrays of custom research electrodes were implanted in the unused space between standard clinical electrodes (clinical monitoring continued uninterrupted during all research procedures). As shown in Figure 2A, one such array consisted of 24 electrodes implanted on the medial face of the occipital lobe in a patient identified by the anonymized subject code YBN. Since the patient was sighted, we used receptive field mapping techniques to identify the regions of the visual field encoded by the cortex underlying each electrode. As expected from the retinotopic organization of visual cortex, there was an orderly organization to these receptive fields (Figure 2B). All were located in the upper right visual field, corresponding to the anatomical location of the electrode array inferior to the calcarine sulcus in the left hemisphere, and more anterior electrodes were associated with more peripheral locations in the visual field. Using the mapped receptive fields, we designed dynamic trajectories of patterned stimulation corresponding to four different letter forms. Successive electrodes in each trajectory were stimulated with small amounts of current (~1 mA) at high frequency (~200 Hz) in rapid temporal sequence (50 ms per electrode). Without any training, the patient was able to make use of the visual percepts created by dynamic current steering and reproduce them on a touchscreen. As shown in Figure 2C, there was a striking correspondence between the predicted and actual letter shape percepts. We tested the subject’s ability to identify different stimulation trajectories using a four alternative forced choice-task. Fifteen of twenty-three presentations were accurately identified, significantly better than chance (66% *vs.* 25%, p = 10^-4^). Similar high accuracy was observed in another patient (subject code YBG) tested with three different stimulation trajectories (77% vs. 33%, p = 10^-6^ from binomial distribution).

**Figure 2.**
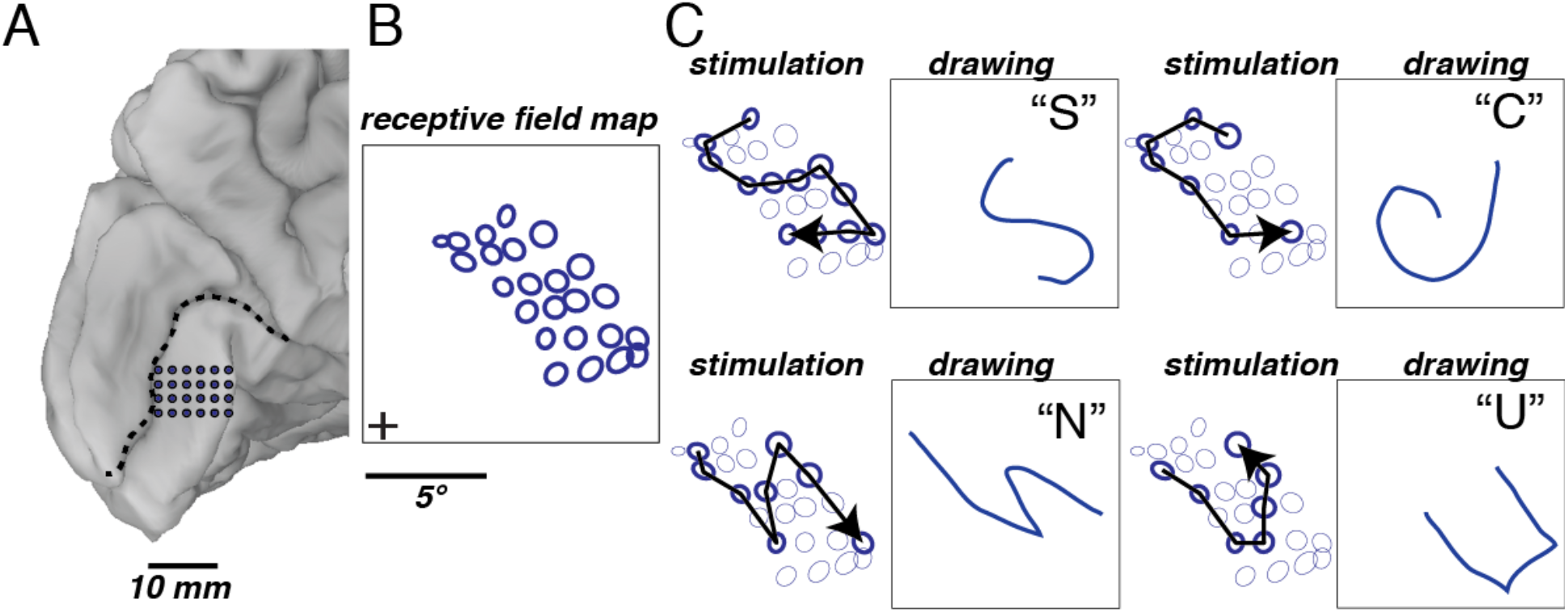
Dynamic current steering guided by visual receptive field mapping. **(A)** Medial view of the left occipital lobe of a sighted patient. Blue circles show the 24 electrodes contained in a grid implanted inferior to the calcarine sulcus (dashed black line). **(B)** To generate receptive field maps, the patient fixated while mapping stimuli were presented. The blue circles show the location in visual space of the mapping stimuli that evoked visual responses for each electrode. **(C)** Dynamic current steering of selected electrodes was used to generate four different artificial percepts. For each percept, left panel shows the stimulated electrodes (bold circles) and direction of temporal sequence of stimulation (arrow). Right panel shows the patient’s drawing of the artificial visual percept and verbal label.

In blind patients, it is not possible to use visual stimuli to map receptive fields in order to design dynamic stimulation trajectories. Therefore, in sighted patient YAY (Figure 3) we tested a different approach in which electrodes were stimulated individually and the patient drew the resulting phosphene on a touch screen. This "phosphene map" allowed for the identification of the visual field region represented by the cortex underlying each electrode (Figure 3B). As expected from the location of the electrodes on the upper bank of the calcarine cortex in the left hemisphere, all phosphenes were in the lower right visual field, with more anterior electrodes producing phosphenes at greater eccentricity. The phosphene map was used to design a dynamic current steering trajectory in the shape of the letter “Z” (Figure 3C). When this stimulation pattern was delivered, the patient was immediately able to accurately reproduce the pattern without any training (Figure 3D; Supplementary Movie S1).

**Figure 3.**
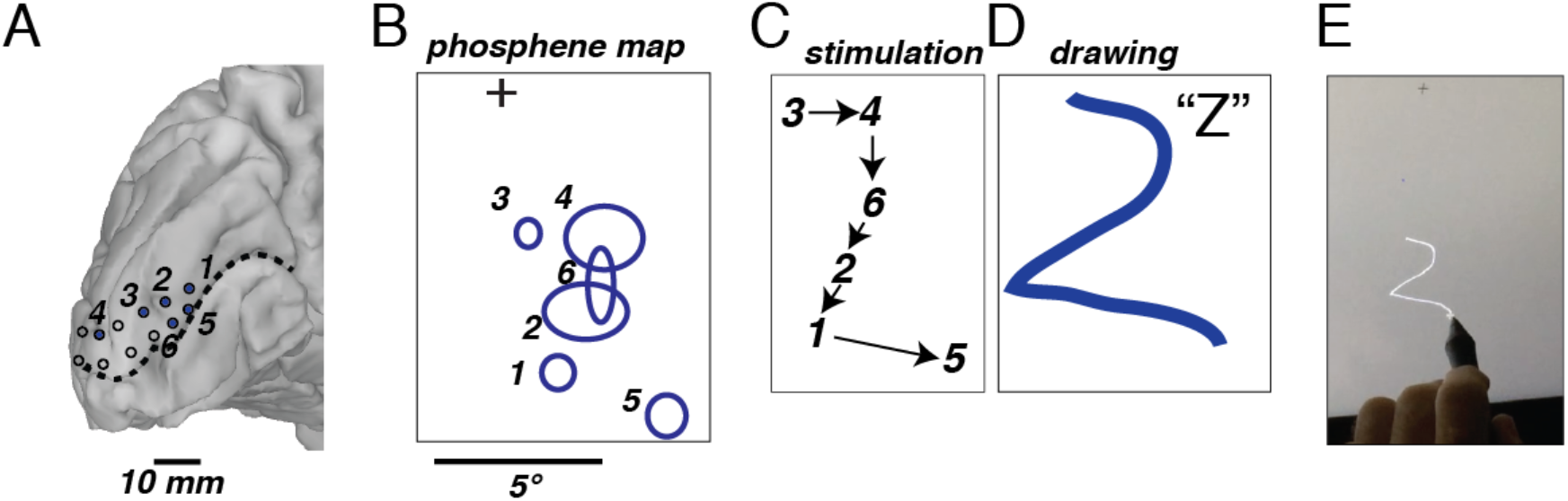
Dynamic current steering guided by phosphene mapping. **(A)** Medial view of the left occipital lobe of a sighted patient. Blue numbered circles show 6 electrodes implanted over the calcarine sulcus (dashed black line) that, when stimulated, created a visual percept. Open circles show electrodes that did not create a visual percept when stimulated. **(B)** The patient fixated while electrical stimulation was delivered to one electrode at a time. The patient drew each phosphene on a touchscreen (bold ellipses, numbered by the corresponding electrode). **(C)** The phosphene map was used to design a stimulation trajectory to produce the artificial percept of the letter “Z”. The black arrows show the temporal sequence of stimulated electrodes. **(D)** The patient drew the artificial percept (blue line) and gave it the verbal label “Z”. **(E)** Still frame from a video of the patient drawing, see *Supplementary Movie 1* for full video.

To determine the utility of the technique in blind individuals, we tested a blind patient with electrodes implanted over visual cortex and controlled through a wireless transmitter (Figure 4). Five of the electrodes produced phosphenes when electrically stimulated, and phosphene mapping revealed two phosphenes located in the upper visual field and three phosphenes located in the lower visual field.

**Figure 4.**
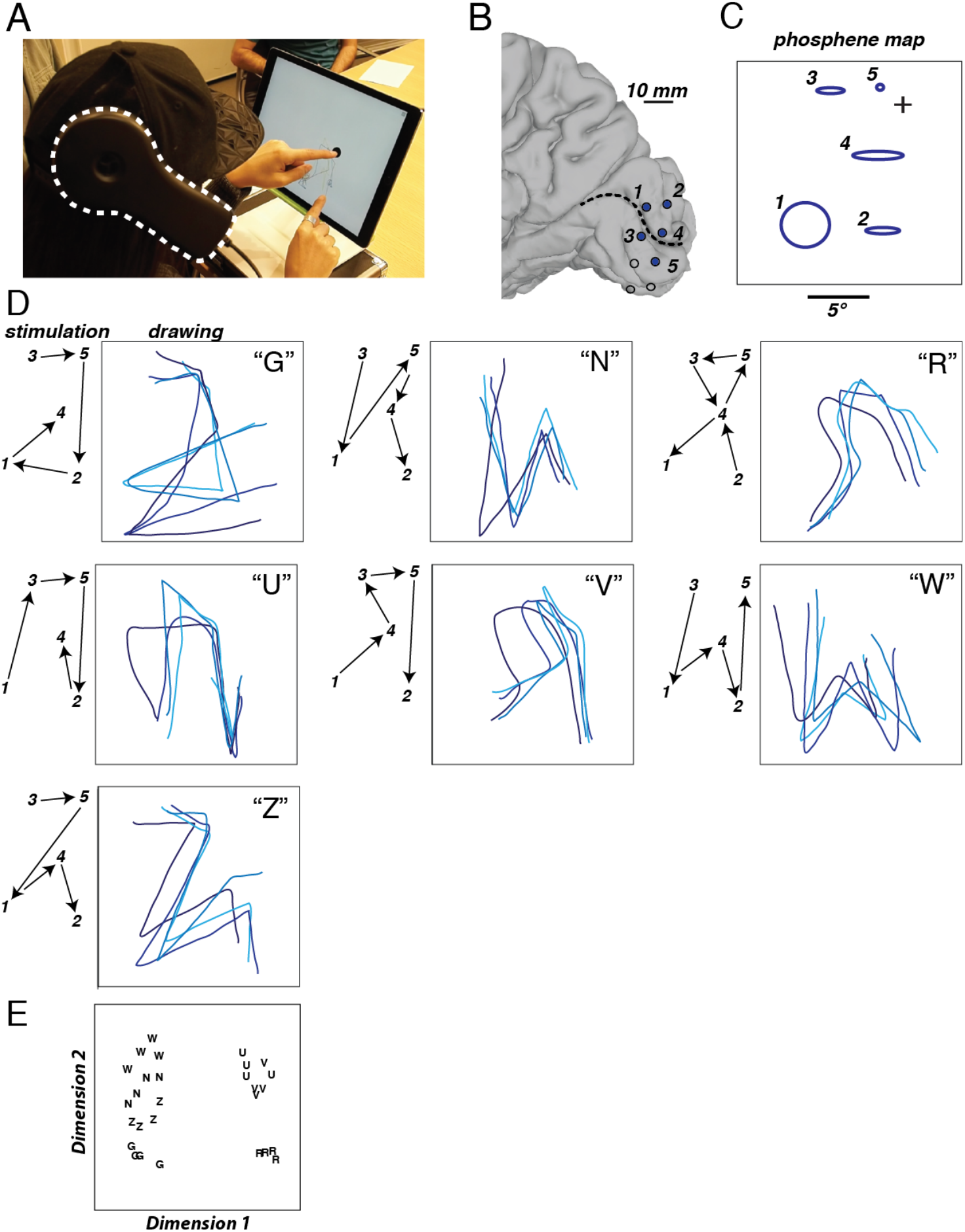
Dynamic stimulation tested in a blind participant. **(A)** The wireless transmitter (dashed white line) was affixed to a cap worn by the participant. The participant placed the index figure of the left hand on a tactile fixation point (black circle) and used the index finger of the right hand to trace the artificial visual percept on the touch screen. Still frame from a video of the patient drawing, see *Supplementary Movie 2* for full video. **(B)** Medial view of a surface model of the participant’s right occipital lobe. Dashed line shows calcarine sulcus, circles show electrode locations. Numbered and filled electrodes were used to create stimulation patterns, open circles show unused electrodes. **(C)** Blue shapes show patient drawing of phosphenes created by stimulation of individual electrodes, numbers corresponding to electrodes in (B). Crosshairs show location of tactile fixation point. **(D)** Seven different letter-like shapes created by seven different dynamic stimulation patterns. Left panel for each shape shows temporal sequence of stimulation (electrodes indicated by numbers connected by black arrows). Right panel for each shape shows patient drawings to different stimulation patterns. Each line illustrates a separate trial (randomly interleaved), colored in different shades of blue for visibility. Letter shows patient mnemonic for the pattern (“Z”; backwards “N”; backwards “R”; upside-down “U”; upside-down “V”; “W”; “Z”). **(E)** Quantification of the drawings produced by the participant for each trial of each stimulation pattern using multidimensional scaling analysis of the pattern drawings. Each letter corresponds to a single trial of the corresponding stimulation pattern.

Seven different dynamic stimulation trajectories were designed to produce letter-like percepts. Without instruction, the blind patient was able to reproduce letter-like shapes that corresponded to the different trajectories (Figure 4D). To assess reliability, the patient was stimulated repeatedly with each of the seven stimulation trajectories (randomly interleaved), drawing the perceived pattern following each trial. The resulting drawings were quantized, correlated, and rendered with multidimensional scaling analysis (Figure 4E).

Repetitions of the same letter-like shape clustered together, while different shapes were centered on different regions of the representational space, demonstrating that the artificial percepts were both distinct and reliable. To further quantify performance, the patient performed a forced-choice discrimination on five of the patterns. Fourteen of fifteen pattern presentations were accurately identified, much higher than expected by chance (93% *vs.* 20%, p = 10^-8^).

The motivation to develop dynamic current steering was provided by repeated failures of the conventional stimulation paradigm to produce shape-like percepts, both in our experience and that of previous investigators (*7, 8*). In one of our failed attempts, 8 electrodes were implanted in the visual cortex of sighted patient YBH (Figure S1). When electrodes were stimulated individually, 8 phosphenes could be reliably mapped. However, when multiple electrodes were stimulated concurrently, shapes were not reported. Instead, only isolated phosphenes were perceived. Typically, the number of individual phosphenes drawn by the patient was fewer than the number of stimulated electrodes, perhaps because spatially adjacent phosphenes coalesced into a single phosphene. When five of the electrodes were stimulated concurrently, the patient reported seeing only two isolated phosphenes (Figure S1C); stimulation of a different set of five electrodes produced a different pattern of two phosphenes (Figure S1D).

**Figure S1.**
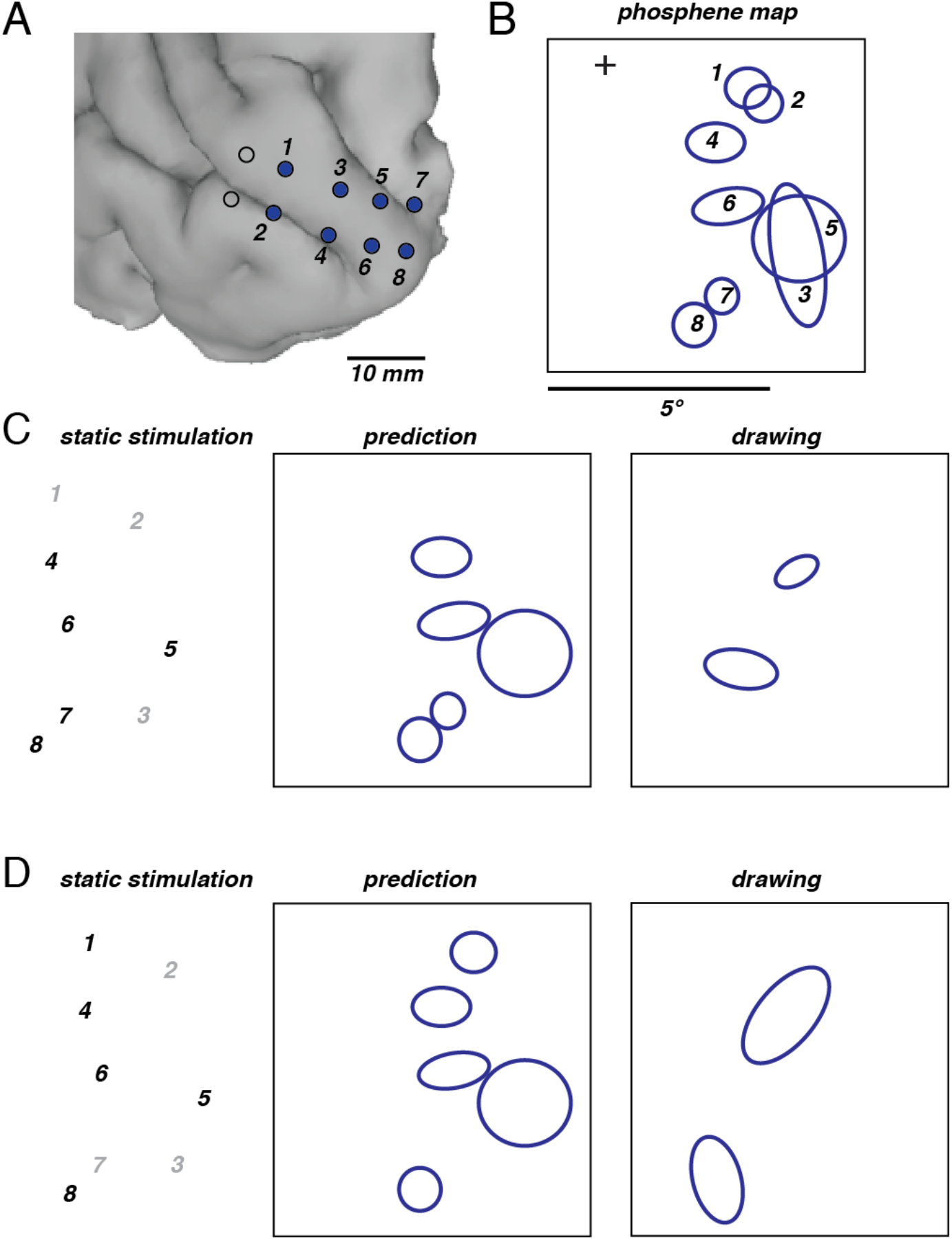
Conventional static stimulation paradigm tested in a sighted subject. **(A)** Posterior lateral view of the subject’s left occipital lobe. Numbered and filled electrodes were stimulated, open circles show non-stimulated electrodes. **(B)** Blue shapes show patient drawing of phosphenes created by stimulation of individual electrodes, numbers corresponding to electrodes in (A). Crosshairs show location of visual fixation point. **(C)** Left panel: bold face numbers show five electrodes stimulated using conventional static stimulation paradigm. Middle panel: percept predicted from phosphene map in (B). Right panel: patient drawing of actual percept. **(D)** Results of conventional static stimulation paradigm for a different set of five electrodes.

## Discussion

Since the 1960s, attempts have been made to restore vision to blind patients with electrical stimulation of the intact visual cerebral cortex (reviewed in *1*). Early attempts were stymied by technological limitations, especially the inability to fabricate miniature electrode arrays and control them with wireless transmitters and receivers. Advances in wireless microelectronics have renewed interest in VCPs, with at least four neuroengineering groups currently developing devices that are either in, or rapidly approaching, the clinical trial stage (*4, 5*). However, these new VCPs rely on the same paradigm for electrical stimulation of visual cortex as the previous generations of VCPs. In the conventional stimulation paradigm, the individual phosphenes created by stimulation of each electrode are treated as pixels in a visual display; turning the pixels on or off is assumed to create a coherent image composed of phosphenes (Chen et al 2009 Vision Research). However, in our experiments, like those of previous attempts (*7, 8, 17*) the conventional stimulation paradigm was never able to evoke useful visual percepts, only isolated phosphenes. In contrast, using the same number of electrodes, the dynamic current steering paradigm was able to evoke a wide variety of patterns that could be immediately reproduced and labeled by the patient without any training.

What is the biology underlying this vast difference in efficacy? Dynamic current steering may be effective because it taps into cortical mechanisms for processing visual motion. In particular, visual motion is perceived for visual stimuli presented without a smooth trajectory, a phenomenon known as apparent motion (*18*). Dynamic current steering evokes cortical activity over a time scale (50 - 100 ms) similar to that which produces apparent motion with real visual stimuli (*19*). Dynamic current steering could leverage the same higher-level cortical areas, such as area MT, that process apparent motion and related motion phenomena, allowing dynamic current steering to evoke a more-or-less natural percept (*20*). As reported by our subjects, a moving phosphene tracing a form is comparable to a real visual motion stimulus, such as a firefly viewed on a dark night.

In contrast, the conventional stimulation paradigm may be ineffective because it evokes patterns of activity very different from those produces by natural visual forms. Natural visual stimuli evoke activity in small populations of tuned neurons (for instance, a “T” presented at the center of gaze will activate foveal neurons in visual cortex selective for vertical and horizontal orientations but not other orientations). In contrast, electrical stimulation of cortex activates neurons representing all orientations, leading to the percept of an amorphous blob. Concurrent stimulation of different locations in the visual cortex produces multiple phosphenes, but the population of activated neurons still includes neurons selective for all orientations, instead of the isolated activity in neurons representing specific orientations that may be a precondition for activating higher cortical regions that are needed for perceiving complex shapes.

Although multiple phosphenes have never been shown to coalesce into a coherent form, they are still capable of conveying information. In the VCP trial of Dobelle and colleagues (*7*), patients were able to learn a correspondence between different combinations of isolated phosphenes and specific letters, a code termed “visual braille.” The visual braille approach has the notable drawback that it requires extensive training to learn a limited set of patterns, restricting the information that can be conveyed to a few arbitrary phosphene-form correspondences learned by the patient. This is in sharp distinction to the ability of the visual system to interpret a never before seen natural scene without training. Like natural vision, dynamic current steering allows the outline of a novel shape to be conveyed through a stimulation trajectory. While we tested only letter-like shapes, the outlines of other common objects, such as faces, houses or cars, could also be traced using the same principles. In combination with modern machine vision algorithms that can rapidly identify objects in visual scenes (*21*), dynamic current steering could be used to give blind participants a rapid outline of salient objects in their environment.

In addition to its ability to convey arbitrary forms, dynamic current steering offers the advantage of delivering much less electrical current to cortical tissue. In the conventional static paradigm, many electrodes are concurrently stimulated, requiring substantial instantaneous power and the possibility of induction of epileptic seizures through synchronized activation. In contrast, in dynamic current steering only a small number of electrodes are concurrently stimulated (two electrodes at a time in our experiments), minimizing instantaneous power requirements. Considering other neural features, such as ongoing oscillations in visual cortex (*22, 23*) could allow further reductions in the amount of current required for dynamic current steering.

Dynamic current steering for VCPs builds on previous efforts to create optimal neural stimulation strategies in human neuroprosthetics. For instance, current steering has been applied both in cochlear implants (*16, 24*) and deep brain stimulators (*25*) although it has not yet found clinical applications in the cerebral cortex.

Advances in technology, including electrical stimulation with high-density grids of electrodes placed on the cortical surface (*26*) or penetrating into the cortex (*27*) and non-electrical stimulation with optogenetic (*28*) magnetothermal (*29*) or focused ultrasound techniques (*30*) promise dramatic increases in our ability to stimulate human cortex. As a general-purpose stimulation paradigm, dynamic current steering can be used in combination with any of these technologies to restore visual function to blind patients.

## Materials and Methods

### Subjects

The subjects for this study consisted of four sighted patients with medically intractable epilepsy (ages 20-54; all male; anonymized subject codes YBN, YAY, YBG and YBH) tested at Baylor College of Medicine (BCM) and one blind patient (age 35; female; anonymized subject code BAA) tested at University of California, Los Angeles (UCLA). All procedures were approved by the Institutional Review Boards of BCM and UCLA.

### Electrodes

Subdural electrodes were implanted on the surface of the occipital lobe. In the epileptic patients, clinical electrodes were implanted for monitoring of epileptogenic activity, with electrode placement guided solely by clinical criteria. Additional research electrodes (embedded in the same silastic strips used for clinical monitoring) were implanted and stimulated for the studies described here. Clinical monitoring continued uninterrupted during experimental sessions.

For patient YBN, the research electrodes consisted of a six by four grid of electrodes (total of 24 electrodes). Each electrode was 0.5 mm in diameter with a center-to-center spacing of 2 mm. For patients YAY, YBG and YBH, the research electrodes consisted of a four by four grid of electrodes (total of 16 electrodes). Each electrode was 0.5 mm in diameter with a center-to-center spacing of 4 mm. For patient BAA, the stimulated electrodes were located on two separate silastic strips. Each strip contained 4 electrodes. Each electrodes was was 3.18 mm in diameter with a center-to-center spacing of 10 mm.

### Electrode Localization

Before surgery, T1-weighted structural magnetic resonance imaging scans were used to create cortical surface models (Figure 1A) with FreeSurfer (*31, 32*) and visualized using SUMA (*33*). Subjects underwent a whole-head CT after the electrode implantation surgery. The post-surgical CT scan and pre-surgical MR scan were aligned using AFNI (*34*) and all electrode positions were marked manually on the structural MR images. Electrode positions were then projected to the nearest node on the cortical surface model using the AFNI program *SurfaceMetrics*.

### Overview of procedures

Epileptic subjects were hospitalized in the epilepsy-monitoring unit for 4 to 14 days after electrode implantation. During all experiments, the patients remained seated comfortably in their hospital bed. A ground pad was adhered to the patient’s thigh, and all electrical stimulation was monopolar. Electrical stimulation currents were generated using a 16-channel system (AlphaLab SnR, Alpha Omega, Alpharetta, GA) controlled by custom code written in MATLAB (Version 2013b, The MathWorks Inc., Natick, MA).

Blind subject BAA acquired blindness at age 27 and had minimal residual light perception. As a component of an early feasibility study for the development of a visual cortical prosthetic, BAA underwent surgical implantation of a neurostimulator normally used to treat epilepsy (RNS System, Neuropace, MountainView, CA). Subject BAA was tested as an outpatient.

### Screening to Determine Responsive Electrodes

We first screened all electrodes implanted on visual cortex to identify responsive electrodes *i.e.* those that produced a phosphene when electrical stimulation was delivered. In each trial, patients verbally reported whether they experienced a localized, brief, visual percept similar to a flash of light. During each trial, an auditory warning tone cued the patients to fixate visual crosshairs. This was followed by a second tone that indicated the beginning of the electrical stimulation period. Electrical stimulation consisted of a train of biphasic pulses (-/+), with 0.1 ms pulse duration per phase, delivered at a frequency of 200 Hz, with an overall stimulus train duration of 200 or 300 ms. Currents tested ranged from 0.3 - 4.0 mA resulting in a total charge delivered of 1.2 – 24 μC per trial. For each electrode, trials were initiated with a low current (0.3-1.0 mA) that gradually increased on successive trials until the patient reported a phosphene. If no phosphene was obtained with a maximum current of 4 mA, then the site was considered unresponsive.

### Quantitative Phosphene Mapping Using Electrical Stimulation

To quantify phosphene locations, additional experiments were performed on each of the electrodes identified in the screening stage. The patient fixated visual crosshairs and electrical stimulation was delivered to a single electrode using the parameters that elicited a phosphene for that electrode in the screening stage. The patient drew the outline of the phosphene on a touchscreen. Multiple trials were typically conducted. On the first trial, the subject was instructed to draw the shape as accurately as possible. On subsequent trials, the patient adjusted the size and location of the phosphene using a custom designed graphical user interface so that it matched the phosphene as precisely as possible. For patient YBH, phosphenes were drawn with a pen and paper instead of a touchscreen. The patient inspected the drawing following the trial. If it did not match the percept, an additional trial was performed and a new drawing created. The paper drawings were digitized using a flatbed scanner. Phosphene drawings for each electrode (touchscreen or pen and paper) were fit with an ellipse for quantification and display. Blind subject BAA by touching and attending to a small Velcro square placed on the screen of an Apple iPad Pro and traced the outline of the phosphene percepts in the appropriate location on the screen of the iPad.

### Receptive Field Mapping Using Visual Stimuli

For sighted patients YBN and YBH, receptive field mapping was performed to measure the visual responses of each electrode (*35*). Patients viewed an LCD screen located approximately 57 cm in front of them. Small black and white flashing checkerboards were presented at different locations on the screen that varied randomly from interval to interval (checkerboard duration 167 ms, blank interval of 167 ms between different locations). To ensure fixation, patients performed a letter detection task at fixation. A 128-channel Cerebus amplifier (Blackrock Microsystems, Salt Lake City, UT) was used to record from the subdural electrodes. An inactive intracranial electrode implanted facing the skull was used as a reference for recording. Signals were amplified, filtered (low-pass: 500 Hz, Butterworth filter with order 4; high-pass: 0.3 Hz, Butterworth filter with order 1) and digitized at 2 kHz. Checkerboard locations that evoked a visual-evoked local field potential were included in the receptive field of the electrode.

### Stimulation Paradigms for Studying Form Vision

For examining form vision, we stimulated multiple electrodes in sequence. Each electrode was stimulated for an identical duration, with a short gap between successive electrodes during which there was no stimulation. For all electrodes, the stimulation frequency was 200 Hz and the pulse width per phase was 100uS. The current amplitude for each electrode was held constant, and was the same used amplitude use for phosphene mapping of individual electrodes (equal to the minimum current that reliably produced a phosphene during the screening stage).

In patient YBN, the current range across the stimulated electrodes was 1.2 mA to 1.5 mA. Each electrode was stimulated for a duration of 50 ms with a no-stimulation interval between stimulation of successive electrodes of 50 ms. In patient YAY, the current range was 0.7-1.5mA (duration 200ms, interval 125 ms). In patient BAA, the current was 2 mA for each electrode (duration 200 ms, interval 2000 ms due to constraints imposed by the electrode control hardware and software). In patient YBH, the current range was 2 – 3 mA (duration 50 ms, no interval between electrodes since all electrodes were stimulated simultaneously).

In patient YBG, the technique of current steering was tested. So-called “virtual electrodes” were created by concurrently stimulating two adjacent electrodes (*16*). A sample stimulation pattern would consist of electrode 1 stimulation at full (100%) current; followed by electrode 1 stimulation at 75% current and electrode 2 stimulation at 25% current, creating a virtual electrode near to electrode 1; followed by electrode 1 stimulation at 50% current and electrode 2 stimulation at 50% current, creating a virtual electrode midway between electrodes 1 and 2; followed by electrode 1 stimulation at 25% and electrode 2 stimulation at 75%, creating a virtual electrode near to electrode 2; followed by electrode 2 stimulation at 100% current. Four real electrodes were used and three virtual electrodes were created between each pair of real electrodes (nine virtual electrodes total) for a total of 13 real and virtual electrodes. The current range across the stimulated electrodes was 1.5 mA to 2.0 mA, with stimulation duration 50 ms and inter-electrode interval 50 ms, resulting in a total sequence duration of 1.3 seconds to traverse all 13 electrodes.

### Behavioral Tests

To assess the subjects’ ability to make perceptual discriminations between different electrical stimulation sequences we used a forced choice discrimination task. Before discrimination testing, the subjects drew the perceived pattern on the touchscreen several times and they were instructed to associate a particular letter or grapheme with each stimulation sequence. During each trial of the discrimination task, a single sequence was presented while the subject fixated, or attended to, a defined place on the touchscreen, and then the subjects gave a verbal report to indicate which of the sequences they had perceived. Sequences were presented in pseudo-random order.

In the blind subject, we also used a multidimensional scaling (MDS) analysis to assess the reliability of differences between the graphemes drawn by the blind subject as a result of different electrical stimulation sequences. One of 7 stimulation sequences was presented on each trial corresponding to one of 7 shapes or graphemes (G, N, R, U, V, W, Z). Each shape was repeated 4 times for a total of 28 trials. For each trial, the drawing made by the subject on the touchscreen was converted into an ordered set of hundreds of evenly spaced circles using Adobe Illustrator. The x and y location of the center of each circle, and hence each point in the original drawing, was then obtained using the regionprops() function in Matlab. So as to have an equal number of points for each drawing, the list of coordinates corresponding to each trial was resampled to obtain exactly 100 points. A correlation matrix 28 x 28 in size was created by obtaining the correlation between the ordered list of x, y points from each trial, and the ordered list of points from every other trial, using the corr2() function in Matlab. The correlation matrix was used as input to Matlab code that performed the MDS analysis.

## Supplementary Materials

Fig. S1. Demonstration of the ineffectiveness of conventional stimulation techniques.

Movie S1: Movie of sighted patient drawing “Z” shape, corresponding to Fig. 3E.

Movie S2: Movie of blind patient drawing upside-down “U” shape, corresponding to Fig. 4A.

## Author contributions

MSB, WB and DY designed and carried out the experiments and wrote the manuscript. PS was responsible for experimental and analysis programming and data collection and analysis. BF contributed to data analysis, data collection, and manuscript preparation. SN and NP were instrumental in the experiments with the blind participant.

## References

1 P. M. Lewis, J. V. Rosenfeld, Electrical stimulation of the brain and the development of cortical visual prostheses: An historical perspective. Brain Res 1630, 208-224 (2016).

2 E. J. Tehovnik, W. M. Slocum, Electrical induction of vision. Neurosci Biobehav Rev 37, 803-818 (2013).

3 W. H. Bosking, M. S. Beauchamp, D. Yoshor, Electrical Stimulation of Visual Cortex: Relevance for the Development of Visual Cortical Prosthetics. Annu Rev Vis Sci 3, 141-166 (2017).

4 A. J. Lowery, in 2013 20th IEEE International Conference on Image Processing. (Melbourne, VIC; Australia, 2013), pp. 1536-1539.

5 P. R. Troyk, in Artificial vision a clinical guide, V. P. Gabel, Ed. (Springer, Cham, Switzerland, 2017), pp. 1 online resource.

6 P. R. Roelfsema, D. Denys, P. C. Klink, Mind Reading and Writing: The Future of Neurotechnology. Trends Cogn Sci 22, 598-610 (2018).

7 W. H. Dobelle, M. G. Mladejovsky, J. R. Evans, T. S. Roberts, J. P. Girvin, “Braille” reading by a blind volunteer by visual cortex stimulation. Nature 259, 111-112 (1976).

8 E. M. Schmidt et al., Feasibility of a visual prosthesis for the blind based on intracortical microstimulation of the visual cortex. Brain 119 (Pt 2), 507-522 (1996).

9 B. A. Wandell, A. M. Rauschecker, J. D. Yeatman, Learning to see words. Annu Rev Psychol 63, 31-53 (2012).

10 T. Hannagan, A. Amedi, L. Cohen, G. Dehaene-Lambertz, S. Dehaene, Origins of the specialization for letters and numbers in ventral occipitotemporal cortex. Trends Cogn Sci 19, 374-382 (2015).

11 W. T. Newsome, E. B. Pare, A selective impairment of motion perception following lesions of the middle temporal visual area (MT). J Neurosci 8, 2201-2211 (1988).

12 R. Gaillard et al., Direct intracranial, FMRI, and lesion evidence for the causal role of left inferotemporal cortex in reading. Neuron 50, 191-204 (2006).

13 A. Kleinschmidt, L. Cohen, The neural bases of prosopagnosia and pure alexia: recent insights from functional neuroimaging. Curr Opin Neurol 19, 386-391 (2006).

14 D. K. Murphey, D. Yoshor, W. H. Bosking, J. H. Maunsell, M. S. Beauchamp, Studying Visual Perception in Human Subjects With fMRI and Intracranial Electrical Stimulation Soc Neurosci Abstr, (2007).

15 H. Keller, J. A. Macy, A. Sullivan, The story of my life. (Doubleday, Page & Company, New York,, 1903), pp. 8 p. l., 3-441 p. incl. facsims.

16 J. B. Firszt, D. B. Koch, M. Downing, L. Litvak, Current steering creates additional pitch percepts in adult cochlear implant recipients. Otol Neurotol 28, 629-636 (2007).

17 W. H. Bosking, B. Foster, P. Sun, M. S. Beauchamp, D. Yoshor, in bioRxiv. (2018).

18 O. Braddick, A short-range process in apparent motion. Vision Res 14, 519-527 (1974).

19 C. Chubb, G. Sperling, Two motion perception mechanisms revealed through distance-driven reversal of apparent motion. Proc Natl Acad Sci U S A 86, 2985-2989 (1989).

20 R. T. Born, D. C. Bradley, Structure and function of visual area MT. Annu Rev Neurosci 28, 157-189 (2005).

21 Y. LeCun, Y. Bengio, G. Hinton, Deep learning. Nature 521, 436-444 (2015).

22 G. Buzsáki, Rhythms of the brain. (Oxford University Press, Oxford; New York, 2006), pp. xiv, 448 p.

23 K. E. Mathewson, G. Gratton, M. Fabiani, D. M. Beck, T. Ro, To see or not to see: prestimulus alpha phase predicts visual awareness. J Neurosci 29, 2725-2732 (2009).

24 R. K. Kalkman, J. J. Briaire, J. H. Frijns, Stimulation strategies and electrode design in computational models of the electrically stimulated cochlea: An overview of existing literature. Network 27, 107-134 (2016).

25 R. E. Gross, M. E. McDougal, Technological advances in the surgical treatment of movement disorders. Curr Neurol Neurosci Rep 13, 371 (2013).

26 D. Khodagholy et al., NeuroGrid: recording action potentials from the surface of the brain. Nat Neurosci 18, 310-315 (2015).

27 P. J. Rousche, R. A. Normann, Chronic recording capability of the Utah Intracortical Electrode Array in cat sensory cortex. J Neurosci Methods 82, 1-15 (1998).

28 K. Deisseroth, Optogenetics: 10 years of microbial opsins in neuroscience. Nat Neurosci 18, 1213-1225 (2015).

29 R. Chen, G. Romero, M. G. Christiansen, A. Mohr, P. Anikeeva, Wireless magnetothermal deep brain stimulation. Science 347, 1477-1480 (2015).

30 W. Legon et al., Transcranial focused ultrasound modulates the activity of primary somatosensory cortex in humans. Nat Neurosci 17, 322-329 (2014).

31 A. M. Dale, B. Fischl, M. I. Sereno, Cortical surface-based analysis. I. Segmentation and surface reconstruction. Neuroimage 9, 179-194 (1999).

32 B. Fischl, M. I. Sereno, A. M. Dale, Cortical surface-based analysis. II: Inflation, flattening, and a surface-based coordinate system. Neuroimage 9, 195-207 (1999).

33 B. D. Argall, Z. S. Saad, M. S. Beauchamp, Simplified intersubject averaging on the cortical surface using SUMA. Human brain mapping 27, 14-27 (2006).

34 R. W. Cox, AFNI: software for analysis and visualization of functional magnetic resonance neuroimages. Computers and biomedical research, an international journal 29, 162-173 (1996).

35 D. Yoshor, W. H. Bosking, G. M. Ghose, J. H. Maunsell, Receptive fields in human visual cortex mapped with surface electrodes. Cereb Cortex 17, 2293-2302 (2007).

